# Effect of E-Cadherin (CDH1) −160C/A polymorphism on prostate cancer risk: a meta-analysis

**DOI:** 10.1101/015123

**Authors:** Yang Bo, He Yi, Wen Xiaofei, Liu Hui, Liao Guoqiang, Liu Feng, Wang Weifeng, Hao Jidong, Ouyang Jun

**Affiliations:** Dept. Urology, Shanghai Zhoupu Hospital, Pudong New District; Dept. Urology, Jiaxing No.1 Hospital, Zhejiang; Dept. Urology, Shanghai East Hospital affiliated to Tongji University; Dept. Urology, Suzhou No.1 Hospital affiliated to Suzhou University

**Author notes:** **Corresponding author** Ouyang Jun, Ph.D, Dept. Urology, Suzhou No.1 Hospital affiliated to Suzhou University. Shizi Street. No 188, Suzhou, Jiangsu Province, China. 215006. These authors contributed equally to this work (Yang Bo, He Yi, Wen Xiaofei).

**Keywords:** E-cadherin (CDH1) polymorphism, homozygotes, heterozygous meta-analysis, prostate cancer

## Abstract

E-Cadherin (CDH1) genetic variations may be involved in invasion and metastasis of various cancers by altering gene transcriptional activity of epithelial cells. However, published studies on the association of CDH1 gene polymorphisms and prostate cancer (PCA)risk remain contradictory, owing to differences in living habits and genetic backgrounds. To derive a more better and comprehensive conclusion, the present meta-analysis was performed. Electronic searches of several databases were conducted for all publications on the association between the CDH1 –160 C/A polymorphism and prostate cancer before Oct 2014. The odds ratio (OR) and its 95% confidence interval (CI) were used for statistical analysis. A total of 7 eligible studies including 1294 cases and 1782 controls were involved in this meta-analysis. Overall, meta-analysis indicated that the −160A allele carriers (AA, CA, AA+CA and A allele) had an increased risk of PCA compared with the homozygotes (CC). In the subgroup analyses by ethnicity, a positive association was found in Asians with A allele, AA, CA, AA+CA genotype and Caucasian descendants with AA genotype, dominant and recessive models. On the contrary, a decreased prostate cancer risk was found in Africans with heterozygous, dominant and allele models. Taken together, this meta-analysis showed that the CDH1 −160A allele might be a risk factor for prostate cancer in Asians and Caucasians. However, this result should be verified by additional population-based studies with large sample sizes.

## Introduction

Prostate cancer is the most common cancer among men and the second leading cause of cancer death in men.^1^ The mechanism of prostatic tumorigenesis is still not fully understood. Evidence from epidemiological and genetic studies provides more focus on the inherited susceptibility to cancer. It is likely that gene environment interactions are involved in tumorigenesis and development.^2^ Among these genetic factors, the E-cadherin (CDH1) gene, consists of a large extracellular domain composed of smaller transmembrane and cytoplasmic domains and five repeat domains.^3^ CDH1, located on chromosome 16q22.1, is one of the most important tumor suppressor genes encoding an adhesion glycoprotein.^4–5^ Several polymorphisms and somatic mutations have been identified in CDH1.^6^ It also plays important roles in such aspects of establishment and maintenance of cell polarity and tissue architecture and intracellular adhesion.^7^ Therefore, abnormal expression of CDH1 is often occurred in a number of human epithelial cancers. Aberrant CDH1 functions have been reported to be associated with malignant transformation of prostatic epithelium as well as metastasis and poor prognosis of PCA.^8–9^ A number of studies had investigated the roles of CDH1 gene polymorphisms in human prostate cancers risk, but the results were not consistent. Therefore, we performed a search of relevant literatures and carried out a meta-analysis to obtain a more accurate evaluation of the association between CDH1 genetic polymorphisms and prostate cancer.

## Methods

### Publication selection

Electronic databases (PubMed, Embase, the Cochrane Controlled Trials Register, the Science Citation Index, and the Chinese Biomedical Database) were searched independently by two authors for all publications regarding the association between the E-Cadherin polymorphism and prostate cancer before Oct 2014. The keywords were as follows: CDH1/ E-Cadherin/ polymorphism/ prostate cancer. A comprehensive search of reference lists of all review articles and original studies retrieved by this method was performed to identify additional reports.. The results were limited to papers with full-text and published in the English language.

### Inclusion and exclusion criteria

Studies estimating the association between CDH1 genetic polymorphisms and prostate cancer risk had to mee all of the following criteria: 1) published in English 2) they were original epidemiological studies on the correlation between CDH1 genetic polymorphisms and prostate cancer susceptibility; 3) case-control studies; 4) sufficien information provided to estimate odds ratios (ORs) with 95% confidence intervals (CIs). However, duplicated studies, case-only studies, case reports, unpublished data letters, comments, and reviews must be excluded.

### Data extraction

Two investigators independently extracted the data and reached a consensus on all the items according to the inclusion criteria listed above. For each study, the following characteristics were collected: author’s first name, year of publication, country of origin, ethnicity, definition of cases, genotyping method, sources of control and case groups, total number of cases and controls, as well as number of cases and controls with AA, CA and CC genotypes.

### Statistical methods

For the control group of each study, the observed genotype frequencies of the CDH1 polymorphism were assessed for Hardy–Weinberg equilibrium using the Pearson chi-squared test; P<0.05 was considered to be statistically significant. Based on both fixed effects and random-effects models, a pooled odds ratio (OR) with 95% confidence interval (CI) was used to assess the strength of association between CDH1 –160 C/A polymorphisms and prostate cancer risk, depending on the heterogeneity of the analysis. The pooled ORs were also assessed for –160 C/A by homozygous (AA vs CC), heterozygous (CA vs CC),recessive (AA vs CC+ CA) and dominant models (AA +CA vs CC) as well as allele comparison (A vs. C).

Heterogeneity was evaluated using the Q test and I^2^ score. If the result of the heterogeneity test was P>0.1, ORs were pooled according to the fixed-effects (Mantel–Haenszel) model. Otherwise, ORs were pooled according to the random-effects (DerSimonian and Laird) model. Sensitivity analysis was performed by omitting one study at a time and recalculating the pooled OR for the remaining studies to assess the stability of the results.

Publication bias was assessed using Egger’s test and Begg’s test. All statistical tests were performed using STATA version 10.0 software (Stata Corporation, College Station, TX, USA). The results were considered statistically significant if the P-value was<0.05.

## Results

### Study Selection

The electronic search strategy identified 31 potentially relevant articles, which were evaluated further in detail, including their titles, abstracts, full text, or a combination of these. Twenty-four articles were excluded (Figure 1). Nine studies were not focused on prostate cancer and seven were not focused on the CDH1 polymorphism. Six studies were laboratory studies, and one study was a systematic review. One study^10^ was eliminated because it did not follow Hardy–Weinberg equilibrium, indicating that these groups might not represent the general population very well. Finally, seven studies^11–17^ on CDH1 genotypes and prostate cancer risk including a total of 1294 prostate cancer cases and 1782 controls were identified.

**Figure 1.**
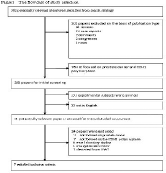
Flow chart of study selection based on inclusion and exclusion criteria.

### Study Characteristics

Table 1 showed the characteristics of the studies included in this meta-analysis. A total of seven publications met the inclusion criteria.^11–17^ All of them are case-control studies. Almost all of the cases were histologically confirmed. The controls were mainly healthy populations except for some having benign prostatic hyperplasia. There were three studies of Asians, two studies of Caucasians, and two studies of Africans. Genotyping methods used in the studies included polymerase chain reaction(PCR)-restriction fragment length polymorphism (RFLP), TaqMan-assay, and dynamic allele-specific hybridization(DASH).The distribution of genotypes among the controls was consistent with the Hardy–Weinberg equilibrium (P>0.05) in all studies.

**Table 1.** Characteristics of Studies Included in the Meta-analysis

| Table1 Characteristics of Studies Included in the Meta-analysis |  |  |  |  |  |  |  |  |  |  |  |  |
| --- | --- | --- | --- | --- | --- | --- | --- | --- | --- | --- | --- | --- |
| Reference | Ethnicity | Definition of cases | Methods | Controls | Genotype |  |  |  |  |  | Case/Control | HWE |
|  |  |  |  |  | Case |  |  | Control |  |  |  |  |
|  |  |  |  |  | AA | CA | CC | AA | CA | CC |  |  |
| Kamoto | Asian | Histologically diagnosed | RFLP | Health BPH | 11 | 71 | 154 | 11 | 85 | 252 | 236/348 | 0.25 |
| Tsukino | Asian | Histologically diagnosed | RFLP | Health | 9 | 77 | 133 | 6 | 66 | 147 | 219/219 | 0.66 |
| Goto | Asian | Histologically diagnosed | RFLP | BPH | 6 | 79 | 115 | 1 | 39 | 119 | 200/159 | 0.25 |
| Bonilla | African | Diagnosed by a pathologist | RFLP | Health | 6 | 50 | 152 | 6 | 67 | 162 | 208/235 | 0.77 |
| Pookot | African | Histologically diagnosed | RFLP | Health | 2 | 9 | 38 | 8 | 46 | 63 | 49/117 | 0.92 |
| Lindstrom | Caucasian | Family positive cases | DASH | Health | 24 | 88 | 87 | 29 | 209 | 268 | 199/506 | 0.15 |
| Hajdinjak | Caucasian | Histologically diagnosed | TanMan | Health | 21 | 72 | 90 | 12 | 81 | 105 | 183/198 | 0.48 |

Abbreviation: HWE, hardy–Weinberg equilibrium.

### Quantitative data synthesis

The results for associations between −160C/A polymorphism and prostate cancer risk and for heterogeneity testing are shown in Table 2. The combined results of all studies showed that significantly increased risk was found between −160C/A polymorphism and prostate cancer (OR 1.84, 95%CI 1.31–2.60 for AA versus CC, P=0.001; OR 1.18, 95% CI 1.01–1.38 for CA versus CC, P=0.04; OR 1.25, 95%CI 1.07–1.45 for the dominant model AA+CA versus CC, P=0.005; OR 1.77, 95%CI 1.27–2.48 for the recessive model AA versus CC + CA, P=0.001, Figure 2)..Further, we detected an association between the −160C/A polymorphism and prostate cancer when examining the comparison of A versus C (OR 1.23, 95%CI 1.04–1.46 for A versus C, P=0.02,Fig 2)

**Table 2.** Meta-analysis of the association between CDH1 −160 C>A polymorphism and prostate cancer risk

| Comparison | Odds<br>ratio | 95% CI | P-value | Heterogeneity |  | Effects<br>model |
| --- | --- | --- | --- | --- | --- | --- |
| | | | | $I^2$ | P-value | |
| A versus C | 1.23 | 1.04-1.46 | 0.02 | 67% | 0.006 | Random |
| Africans | 0.65 | 0.44-0.97 | 0.03 | 66% | 0.08 |  |
| Asians | 1.55 | 1.23-1.95 | 0.001 | 15% | 0.31 |  |
| Caucasians | 1.22 | 0.87-1.72 | 0.25 | 0% | 0.80 |  |
| AA versus CC | 1.84 | 1.31-2.60 | 0.001 | 12% | 0.34 | Fixed |
| Africans | 0.74 | 0.30-1.84 | 0.52 | 0% | 0.35 |  |
| Asians | 1.95 | 1.05-3.64 | 0.04 | 0% | 0.50 |  |
| Caucasians | 2.33 | 1.45-3.73 | 0.001 | 0% | 0.65 |  |
| CA versus CC | 1.18 | 1.01-1.38 | 0.04 | 70% | 0.003 | Random |
| Africans | 0.64 | 0.44-0.94 | 0.02 | 72% | 0.06 |  |
| Asians | 1.50 | 1.19-1.90 | 0.001 | 29% | 0.25 |  |
| Caucasians | 1.19 | 0.91-1.55 | 0.21 | 0% | 0.42 |  |
| AA+CA versus CC | 1.25 | 1.07-1.45 | 0.005 | 73% | 0.001 | Random |
| Africans | 0.66 | 0.46-0.94 | 0.02 | 75% | 0.05 |  |
| Asians | 1.54 | 1.23-1.93 | 0.001 | 38% | 0.20 |  |
| Caucasians | 1.33 | 1.03-1.71 | 0.03 | 0% | 0.41 |  |
| AA versus CC+CA | 1.77 | 1.27-2.48 | 0.001 | 0% | 0.62 | Fixed |
| Africans | 0.88 | 0.36-2.19 | 0.79 | 0% | 0.50 |  |
| Asians | 1.75 | 0.94-3.25 | 0.08 | 0% | 0.58 |  |
| Caucasians | 2.15 | 1.37-3.39 | 0.001 | 0% | 0.81 |  |

Abbreviation: CI, confidence interval

**Figure 2.**
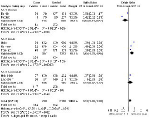
Forest plot showing the association of the CDH1 −160C>A A allele with risk of prostate cancer compared with the C allele. The squares and horizontal lines correspond to the study-specific OR and 95% CI. The diamond represents the summary OR and 95% CI. Abbreviations: CI, confidence interval; OR, odds ratio.

Ethnicity subgroup analysis revealed that −160C/A polymorphism was related with increased risk of prostate cancer in Asian when examining the comparison of A versus C (OR 1.55, 95%CI 1.23–1.95, P=0.001), AA versus CC (OR 1.95, 95%CI 1.05–3.64, P=0.04,),CA versus CC (OR 1.50, 95%CI 1.19–1.90, P=0.001) and AA+CA versus CC (OR 1.54, 95% CI 1.23–1.93, P=0.001, Figure 3). However, no significant difference was found in the recessive model (AA versus CC+CA, OR 1.75, 95%CI 0.94–3.25, P=0.08)

**Figure 3.**
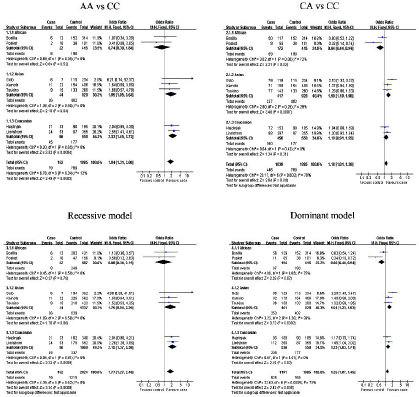
Forest plot describing the association of the CDH1 −160C>A genetic models (AA versus cc, CA versus cc, AA + CA versus CC, AA versus CC + CA) with the risk of prostate cancer. Abbreviations: CI, confidence interval; OR, odds ratio.

In the Caucasian group, there was a significant increased risk between −160C/A polymorphism and prostate cancer risk when examining the comparison of AA versus CC (OR 2.33, 95%CI 1.45–3.73, P=0.001), AA+CA versus CC (OR 1.33, 95% CI 1.03–1.71, P=0.03) and AA versus CC+CA(OR 2.15, 95%CI 1.37–3.39, P=0.001, Figure 3). However, no significant difference was found in the other genotype distributions (A versus C: OR 1.22, 95%CI 0.87–1.72, P=0.25; CA versus CC: OR 1.19, 95%CI 0.91–1.55, P=0.21)

In the Afican group, results indicated that −160C/A polymorphism was related with decreased prostate cancer risk when examining the comparison of A versus C (OR 0.65, 95%CI 0.44–0.97, P=0.03),CA versus CC (OR 0.64, 95%CI 0.44–0.94, P=0.02) and AA+CA versus CC (OR 0.66, 95%CI 0.46–0.94, P=0.02, Figure 3).However, no significant association was found in any genetic models (AA versus CC, OR 0.74, 95%CI 0.30–1.84, P=0.52; AA versus CC+CA, OR 0.88, 95%CI 0.36-2.19, P=0.79).

### Tests of heterogeneity

Statistically significant heterogeneity was found between the trials using the Q statistic and I^2^score (A versus C, P=0.006, I^2^=67%; CA versus CC, P=0.003, I^2^=70%; the dominant model AA+CA versus CC, P=0.001, I^2^=73%). The random-effects model was employed in these studies. There was no significant heterogeneity between the following comparisons: AA versus CC (P=0.34, I^2^=12%) and recessive model AA versus CC+CA (P=0.62, I^2^=0%, Table 2), and the fixed-effects model was employed in these studies.

### Sensitivity analysis

Sensitivity analysis was conducted to assess the stability of the results by sequential removal of each eligible study. The significance of the pooled ORs was not influenced by any single study, indicating that our results were statistically robust(data not shown).

### Publication bias

Begg’s test and Egger’s test were used to assess for publication bias. Egger’s weighted regression did not indicate publication bias (A versus C, P=0.855; AA versus CC, P=0.942; CA versus CC, P=0.832; dominant model AA + CA versus CC, P=0.686; recessive model AA versus CC + CA, P=0.629).This was confirmed by Begg’s rank correlation (A versus C, P=1.000; AA versus CC, P=1.000; CA versus CC, P=0.443; dominant model AA + CA versus CC, P=0.443; recessive model AA versus CC + CA, P=1.000, Table 3).

**Table 3.** Publication bias test for CDH1 −160C>A polymorphism

| Comparison |  |  | Egger's test |  |  | Begger's test<br>P value |
| --- | --- | --- | --- | --- | --- | --- |
|  |  |  | Coefficient | P value | 95%CI |  |
| A | versus | C | 0.332 | 0.855 | -6.594, 5.608 | 1.000 |
| AA | versus | CC | 0.425 | 0.942 | -4.721, 4.225 | 1.000 |
| CA | versus | CC | -0.573 | 0.832 | -4.273, 3.138 | 0.443 |
| AA+CA | versus | CC | -0.211 | 0.686 | -5.286, 4.257 | 0.443 |
| AA | versus | CC+AC | 0.813 | 0.629 | -4.672, 5.142 | 1.000 |

Abbreviation:CI,confidence interval

## Discussion

Many studies have attempted to reveal the genetic basis of prostate cancer. Despite suggestive evidence of gene association, reports have been difficult to replicate, indicating that prostate cancer is more genetically heterogeneous than initially believed. The polymorphisms of CDH1 may play a critical role in the tumorigenesis, development and prognosis of sever kinds of cancer, such as urothelial cancer, colorectal cancer, gastric cancer, prostate cancer, breast cancer and esophageal cancer.^18–23^ Because PCA is one of the most common malignant diseases among men and a number of studies have reported a function of the CDH1 −160 C>A polymorphism in PCA risk with inconclusive results, we performed this meta-analysis to estimate the association specifically. At the same time, because the same polymorphism seemed to have different functions in cancer susceptibility among different ethnic populations and because the frequencies of single-nucleotide polymorphisms might be different among different ethnic groups, subgroup analyses on the basis of ethnicity were conducted.

We concluded that rs16260 A allele was obviously associated with increased cancer risk based on 1294 cases and 1782 controls in overall pooled results from 7 studies. A stratified analysis by cancer type indicated that rs16260 AA genotype increased risk of prostate cancer. Subsequently stratified analysis by country indicated borderline increased cancer risk was found in Asian and Caucasian population. In contrast, results appeared in African population revealed that −160C/A polymorphism was related with decreased risk of prostate cancer. This may be related with different genetic background and the environment exposure^24^ or the different frequency of rs16260 A allele variant in this study. In addition, it is also possible that the observed ethnic differences may be due to chance because studies with small sample size may have insufficient statistical power to detect a slight effect or may have gene-rated a fluctuated risk estimate.^25^ Considering the limited studies and population numbers of Africans included in the meta-analysis, our results should be interpreted with caution

Heterogeneity is a potential problem when interpreting the results of all meta-analyses.^26^ Significant between-study heterogeneity existed in overall comparisons. After subgroup analyses by ethnicity, the heterogeneity was effectively decreased or removed in Europeans and Asians. The reason might be that differences of genetic backgrounds and the environment existed among different ethnicities.

Some limitations of this meta-analysis should be acknowledged. First, the influence of the genetic variant may be masked by the presence of other as yet unidentified genes involved in carcinogenesis, which restricted our evaluation of potential gene–gene interactions. Secondly, while publication bias was not detected in three polymorphisms of CDH1, publication bias which we did not detect might also exist in other polymorphisms owing to small amount of studies. Thirdly, controls were not uniformly defined. Although the healthy populations were the main source of the controls, some of them might be patients. Fourthly, in the subgroup analysis, the number of cases and controls was relatively small in different cancers, races and source of controls, not having sufficient statistical power to achieve the real association. At last, our results were based on unadjusted evaluation, so a more precise analysis should be conducted with adjustment for other variables, eg, environmental factors. Larger and better designed studies are needed to evaluate further the association between the CDH1 polymorphism and prostate cancer risk, including considering the possibility of gene–gene or SNP–SNP interactions and the possibility of linkage disequilibrium between polymorphisms.

In conclusion, it is worthwhile searching for polymorphic variants influencing the risk of prostate cancer. This meta-analysis provides evidence of an association between CHD1 –160 C>A polymorphism and prostate cancer risk, supporting the hypothesis that the CHD1 –160 C>A A allele may act as a risk factor for prostate cancer in Asians but not in Caucasians. However, our results should be interpreted with caution because of some limitations. Given that the results of this meta-analysis are preliminary and may be biased by the relatively small number of subjects, additional population-based studies including large sample sizes should be conducted to verify the association of CDH1 polymorphism in prostate cancer.

## Acknowledgements

This study was funded in part by academic pacesetter program of Shanghai Pudong New District Health System. (PDRd2011-08). Research project of Shanghai health bureau (2013-609). Natural Science Foundation of China (81070591).

**Competing interests** None

